# Design and Synthesis of Peptide-Polyester Conjugates for Cell-Mediated Scaffold Degradation

**DOI:** 10.1101/2025.10.06.680674

**Authors:** Korina Vida G. Sinad, Natasha K. Hunt, Srujan Singh, Kelly B. Seims, Yingjie Wu, E. Thomas Pashuck, Warren L. Grayson, Lesley W. Chow

## Abstract

Biodegradable thermoplastic polyesters are promising biomaterials for tissue engineering due to their processability and mechanical properties. Polycaprolactone (PCL) is particularly attractive for load-bearing applications but does not degrade at the same rate as new tissue formation, which may compromise functional regeneration. This study presents a strategy for cell-mediated scaffold remodeling by incorporating a protease-cleavable peptide directly into the PCL backbone. Linear peptide–PCL conjugates were synthesized with poly(ethylene glycol) (PEG) spacers flanking the peptide to enhance protease access. A functional proteomics approach was used to identify a fast-degrading peptide sequence (Fast) selectively cleaved by multiple cell types. Conjugates containing Fast or its scrambled control (ScrFast) were combined with an RGDS-PCL conjugate and fabricated into scaffolds. Including Fast and ScrFast peptides did not impair cell adhesion to the scaffolds. Cy3 labeling enabled real-time quantification of scaffold degradation in the presence of collagenase or human mesenchymal stromal cells (hMSCs). Fast-PCL scaffolds degraded significantly faster than ScrFast-PCL in both conditions, demonstrating sequence-dependent and cell-directed resorption. Integrating protease-sensitive peptides into the polymer backbone is therefore an effective approach to fabricate solid scaffolds that degrade in response to cells. This platform can be adapted to couple cellular processes to scaffold remodeling to enhance tissue regeneration.

## 1. Introduction

Biomaterials for tissue engineering (TE) are designed to replace or restore the function of damaged or degenerated tissues by providing temporary physical support for cellular infiltration and tissue regeneration, while gradually being resorbed as new tissue forms.^[1–5]^ Biodegradable thermoplastic polyesters like polycaprolactone (PCL), poly(lactic acid) (PLA), poly(glycolic acid) (PGA), and poly(lactide-*co*-glycolic acid) (PLGA) polymers are attractive scaffold materials due to their ability to degrade under physiological conditions.^[6,7]^ Moreover, they offer key benefits, such as batch-to-batch reproducibility, controllable chemistries and microstructures, and clinical use in FDA-approved products.^[4,8–10]^ They also possess mechanical properties suitable for orthopedic applications and can be easily processed into patient-specific geometries for complex defect repair.^[11–15]^ For example, PCL 3D printed with decellularized bone into anatomically shaped scaffolds and implanted with human adipose-derived stem cells significantly enhanced craniofacial bone healing in critical-sized defects.^[16,17]^ While PCL-based scaffolds provide adequate mechanical support, they demonstrate minimal degradation *in vivo* (∼24 – 48 months) that contrasts sharply with the typical timeline of bone healing (∼6 months).^[6,16,18–21]^ Solid scaffolds that degrade too slowly may hinder tissue growth or leave voids once they degrade in the injury site post-healing, compromising long-term success.^[5,16,21,22]^ Conversely, those that degrade too rapidly risk losing mechanical integrity before sufficient tissue has formed.^[23,24]^

Scaffold degradation can be controlled through factors that affect polymer degradation rate, such as modifying functional groups, copolymer composition, and microstructure.^[24,25]^ PCL degradation can be accelerated by copolymerizing with faster-degrading polymers like PLA or PGA, but this approach results in scaffolds with pre-set degradation rates.^[26,27]^ This limits adaptability to patient-specific healing dynamics, which are influenced by local factors (e.g., blood supply and infection), systemic factors (e.g., age, nutrition, and menopause), or mechanical factors (e.g., stress and movement).^[28–36]^ These complexities motivate the need for ‘smart’ scaffolds that respond to cellular activity to match scaffold degradation to tissue remodeling and regeneration processes.

Biological events, such as protease activity, have been leveraged to induce cell-mediated biomaterial degradation.^[37–40]^ For example, cells secrete proteases like matrix metalloproteinases (MMPs) and the serine protease plasmin, to remodel their surrounding extracellular matrix (ECM) during migration and proliferation.^[41–43]^ Synthetic hydrogel systems, particularly those composed of poly(ethylene glycol) (PEG), have been crosslinked with protease-sensitive peptides for tunable degradation in response to cell-secreted enzymes.^[44–47]^ Degradation rate can be controlled by changing the number of proteolytic cleavage sites and incorporating peptide sequences with varying protease specificity.^[41,48]^ Hydrogels have been the dominant material in this space due to their water-swollen architecture that increases access to the protease-sensitive peptides. However, their application is typically limited to drug delivery, injectable therapeutics, and soft tissue regeneration because of their relatively fast degradation rate (e.g., days to weeks) and low mechanical strength. ^[49–54]^

Protease-sensitive degradation has recently been expanded to more mechanically robust materials, such as solid thermoplastic polymers including polyurethanes and polyesters.^[55–57]^ Fung and colleagues synthesized a linear poly(ester-arylate) containing peptides with protease-specific sensitivity.^[55]^ Incorporating peptides within the polymer backbone resulted in protease-specific polymer surface resorption, demonstrating the potential for employing protease-guided degradation in hydrophobic thermoplastic polymers.^[55]^ Similarly, elastase-sensitive and collagenase-sensitive sequences have been incorporated into polyurethane and poly(ether ester) systems, often aided by PEG to enhance hydrophilicity and protease accessibility.^[58–61]^ These studies underscore the potential for integrating protease-responsive motifs into biodegradable thermoplastics to create solid scaffolds that degrade in concert with cellular activity.

Here, we developed a proteomics-guided strategy to create solid polymeric scaffolds that undergo selective, cell-driven remodeling. Unlike prior systems that relied on known peptide substrates, we used a functional proteomic workflow to identify a peptide sequence that is cleaved by multiple cell types. We designed a robust synthetic approach that integrates the peptide within the backbone of PCL to create linear peptide-PCL conjugates with PEG spacers flanking the peptide to facilitate protease access **(Figure 1)**. Scaffolds fabricated with these conjugates exhibited sequence-specific enzymatic cleavage in collagenase and in the presence of cells. This work introduces a versatile platform that combines proteomics-driven peptide discovery with polymer conjugate design to create mechanically robust materials in which resorption is dictated by cellular activity.

**Figure 1.**
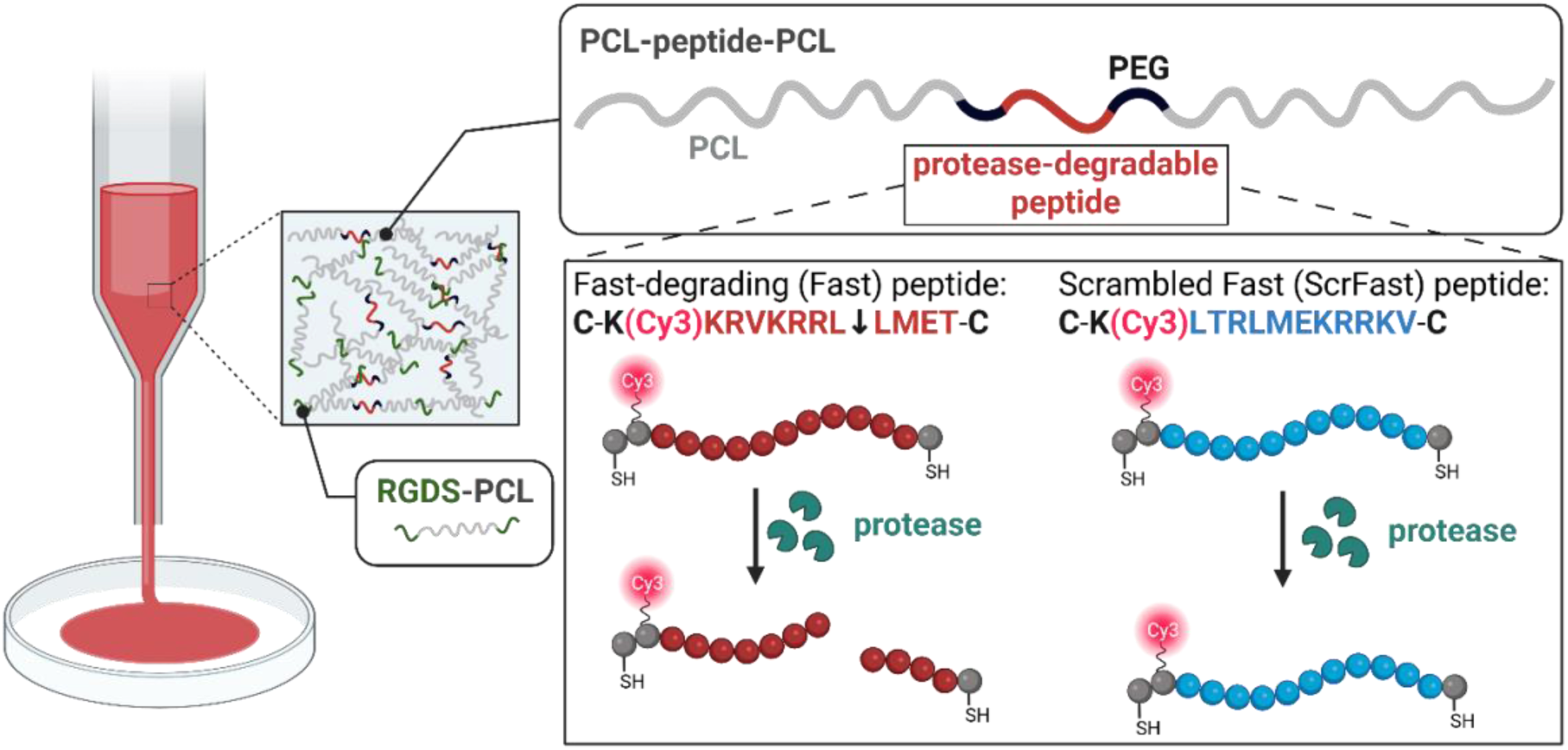
Schematic overview of biomaterial design. A fast-degrading peptide (Fast) and the scrambled control (ScrFast) were integrated into the backbone of a maleimide-functionalized polymer via Michael addition to synthesize Fast-PCL and ScrFast-PCL conjugates. Scaffolds were prepared by solvent-casting the conjugates. An RGDS-PCL conjugate was included for cell adhesion. Protease-mediated scaffold degradation was quantified using fluorescence measurements of solutions collected from scaffolds incubated in protease-containing media or in the presence of cells. Created with BioRender.com.

## 2. Results and Discussion

### 2.1. Design and Synthesis of Protease-Sensitive Peptide-Polymer Conjugate

We applied a functional proteomics-based approach to systematically identify novel protease-cleavable peptides with broad relevance across multiple cell types.^[62,63]^ This method captures the cumulative enzymatic activity of both soluble and membrane-bound proteases secreted by human cell types, allowing for the discovery of peptide substrates that are more biologically representative than canonical sequences like GPQGIWGQ (PanMMP).^[62,64]^ Within this peptide library, we selected a peptide sequence, KRVKRRLLMET (Fast), that exhibited rapid proteolytic cleavage after 24 hours by human mesenchymal stromal cells (hMSCs), fibroblasts, and different macrophage phenotypes compared to KGPQGIWGQK containing the PanMMP sequence and PPDKTSPEPA **(Figure 2)**. The Fast peptide therefore provided as a model substrate to demonstrate the feasibility of engineering solid scaffolds capable of cell-driven, sequence-specific remodeling.

**Figure 2.**
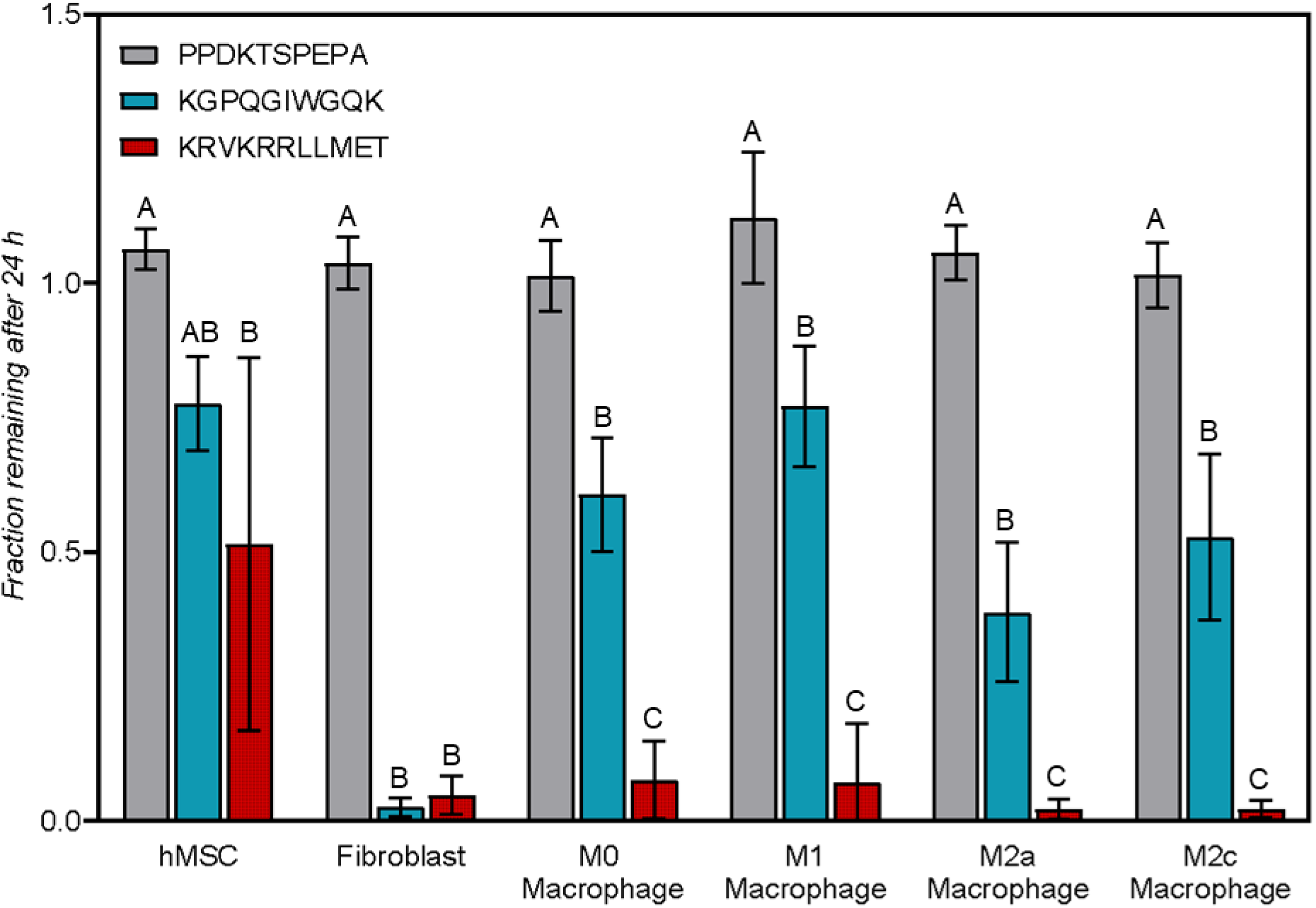
Fractions of candidate peptides remaining after 24 hours of exposure to fresh media with hMSCs, fibroblasts, or different macrophage phenotypes. KRVKRRLLMET generally demonstrated significant degradation compared to PPDKTSPEPA and KGPQGIWGQK. Different letters within a group indicate statistically significant differences (p < 0.05).

We synthesized a scrambled version of the Fast peptide, LTRLMEKRRKV (ScrFast), in which the positions of uncharged amino acids (L, M, V) were swapped with charged amino acids (E, K, R) to ensure the physiochemical properties of the scrambled peptide were significantly different. In contrast to Fast, ScrFast exhibited negligible cleavage when exposed to hMSCs, fibroblasts, and multiple macrophage phenotypes for 24 hours **(Figure 3)**. These results confirmed that degradation was driven by precise protease recognition rather than general peptide susceptibility. ScrFast was used as the negative control in subsequent scaffold studies.

**Figure 3.**
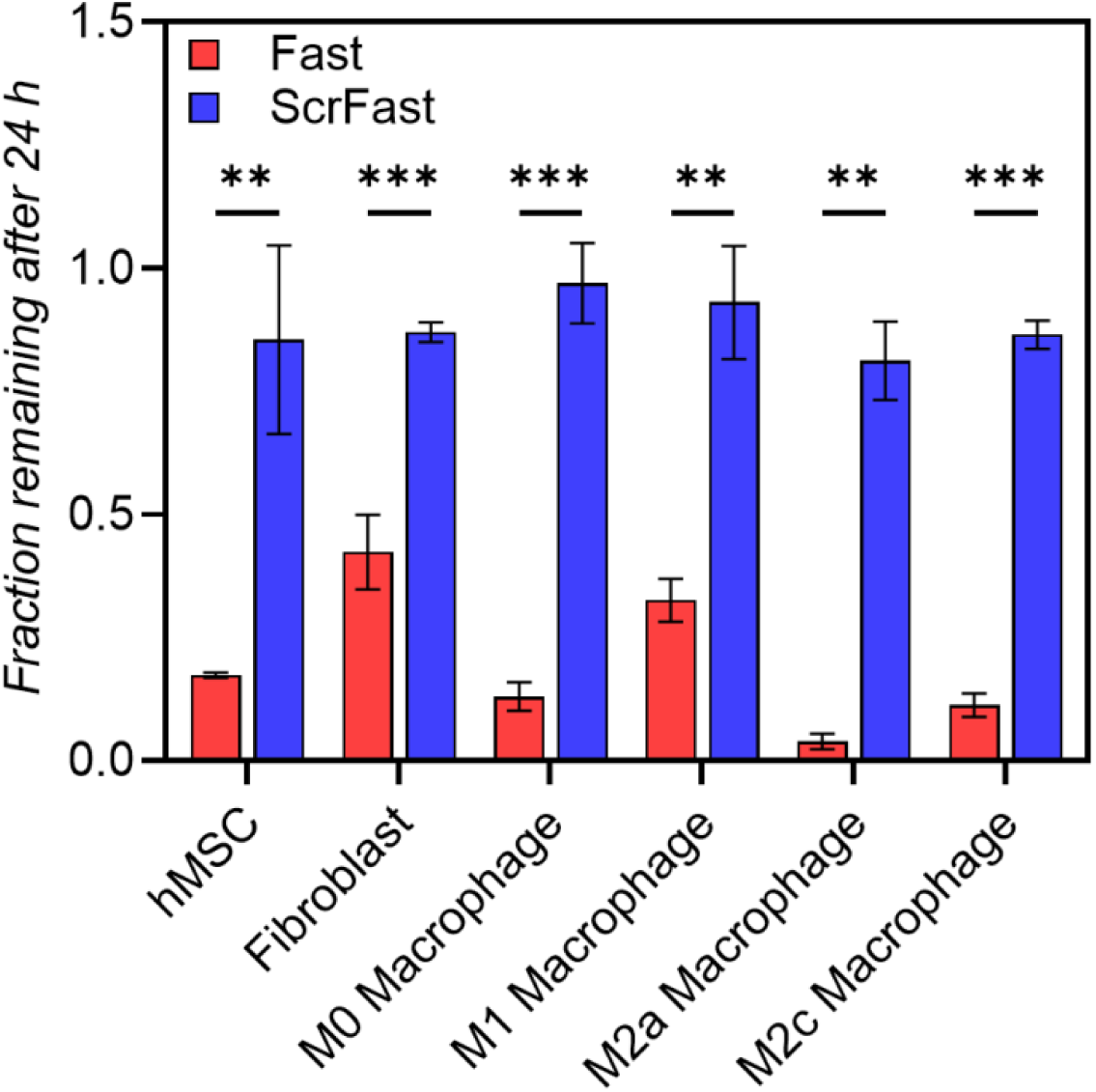
Fractions of KRVKRRLLMET (Fast) and LTRLMEKRRKVK (ScrFast) peptides remaining after 24 hours of exposure to fresh media with either hMSCs, fibroblasts, or macrophages. The Fast peptide showed significant degradation in the presence of all cell types compared to ScrFast peptide (N=3; *p < 0.05, **p < 0.01, ***p < 0.001).

We integrated the Fast and ScrFast peptides into a PCL conjugate backbone to compare sequence-specific protease-mediated scaffold degradation. Fast and ScrFast peptides were modified with a cysteine group on both termini to allow facile conjugation to polymers following different chemistries, including thiol-Michael additions and thiol-ene/thiol-yne radical-mediated reactions.^[65]^ A water-soluble Cy3 fluorophore was added to the peptide to monitor degradation via release of fluorescence into surrounding media. Successful synthesis and purification of the modified Fast and ScrFast peptides were confirmed via high performance liquid chromatography (HPLC), matrix-assisted laser desorption/ionization time-of-flight mass spectrometry (MALDI-ToF MS), and proton nuclear magnetic resonance spectroscopy (^1^H NMR) **(Figures S1 and S2)**. Cy3-modified Fast and ScrFast peptides were coupled to maleimide-functionalized PCL to create linear peptide-PCL conjugates, Fast-PCL and ScrFast-PCL, respectively **(Figure S3)**. Incorporating peptides into the backbone of hydrophobic polymers like PCL can reduce proteolytic access. Proteolytic cleavage of peptides requires access to the active site of enzymes.^[66,67]^ We therefore flanked the peptides with flexible, hydrophilic PEG spacers to increase steric accessibility of the protease-substrate peptides.^[58–60]^ Successful synthesis of maleimide-functionalized PCL and incorporation of the peptides into the PCL backbone was confirmed by ^1^H NMR (**Figures S4 and 4)**. Conjugation efficiency was 88% for Fast-PCL and 82% for ScrFast-PCL based on integration at 7.71 ppm attributed to the indole -C_7_-H (m, 2H, b) and the peak at 4.07 ppm (m, 912H, 2) assigned to -O-CH_2_-of the PCL backbone **(Figure 4)**.

**Figure 4.**
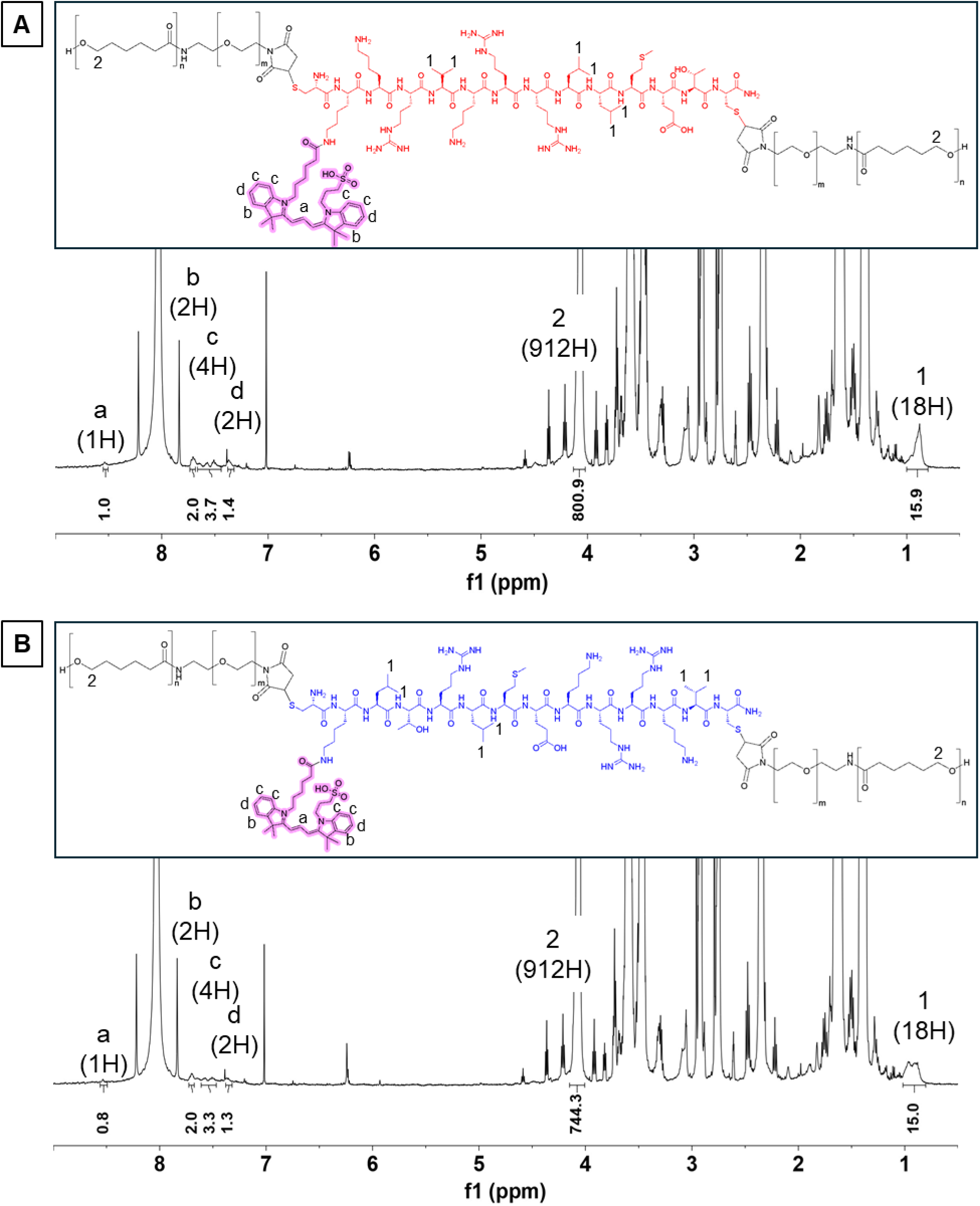
Representative ^1^H NMR (500 MHz, DMF-d_7_) and corresponding chemical structures of synthesized peptide-PCL conjugates: (A) Fast-PCL and B) ScrFast-PCL: δ = 8.54 (m, 1H, vinyl H, a), 7.71 (m, 2H, Ar-H, b), 7.54 (m, 4H, Ar-H, c), 7.37 (m, 2H, Ar-H, d), 4.07 (t, 912H, -O-CH_2_-, 2), and 0.89 (m, 18H, -CH_3_, 1).

Scaffold disks were successfully fabricated by solvent-casting inks containing Fast-PCL and ScrFast-PCL conjugates onto glass petri dishes. We included an RGDS-PCL conjugate in the ink prior to casting to support cell adhesion. We previously showed that RGDS-PCL can be incorporated prior to scaffold fabrication to functionalize the scaffold surface and promote cell adhesion.^[68]^ RGDS and RGDS-PCL were successfully synthesized and characterized **(Figures S5 and S6)**.^[68]^ Both scaffolds appeared bright pink due to the Cy3 fluorophore attached to the peptides **(Figure 5A)**. Representative SEM images revealed similar surface morphologies between Fast-PCL and ScrFast-PCL scaffolds, indicating that peptide incorporation did not noticeably alter scaffold topography. **(Figure 5B)**. Fast-PCL and ScrFast-PCL scaffolds seeded with hMSCs and stained with Hoechst 33342 showed cell nuclei present on both scaffolds at Day 4 **(Figure 5C)**. DNA quantification indicated no significant differences in the number of cells adhered to each scaffold **(Figure 5D)**. These results demonstrated that Fast-PCL and ScrFast-PCL could be successfully fabricated into similar scaffolds and that integrating these peptides into the conjugate backbone did not negatively impact cell adhesion.

**Figure 5.**
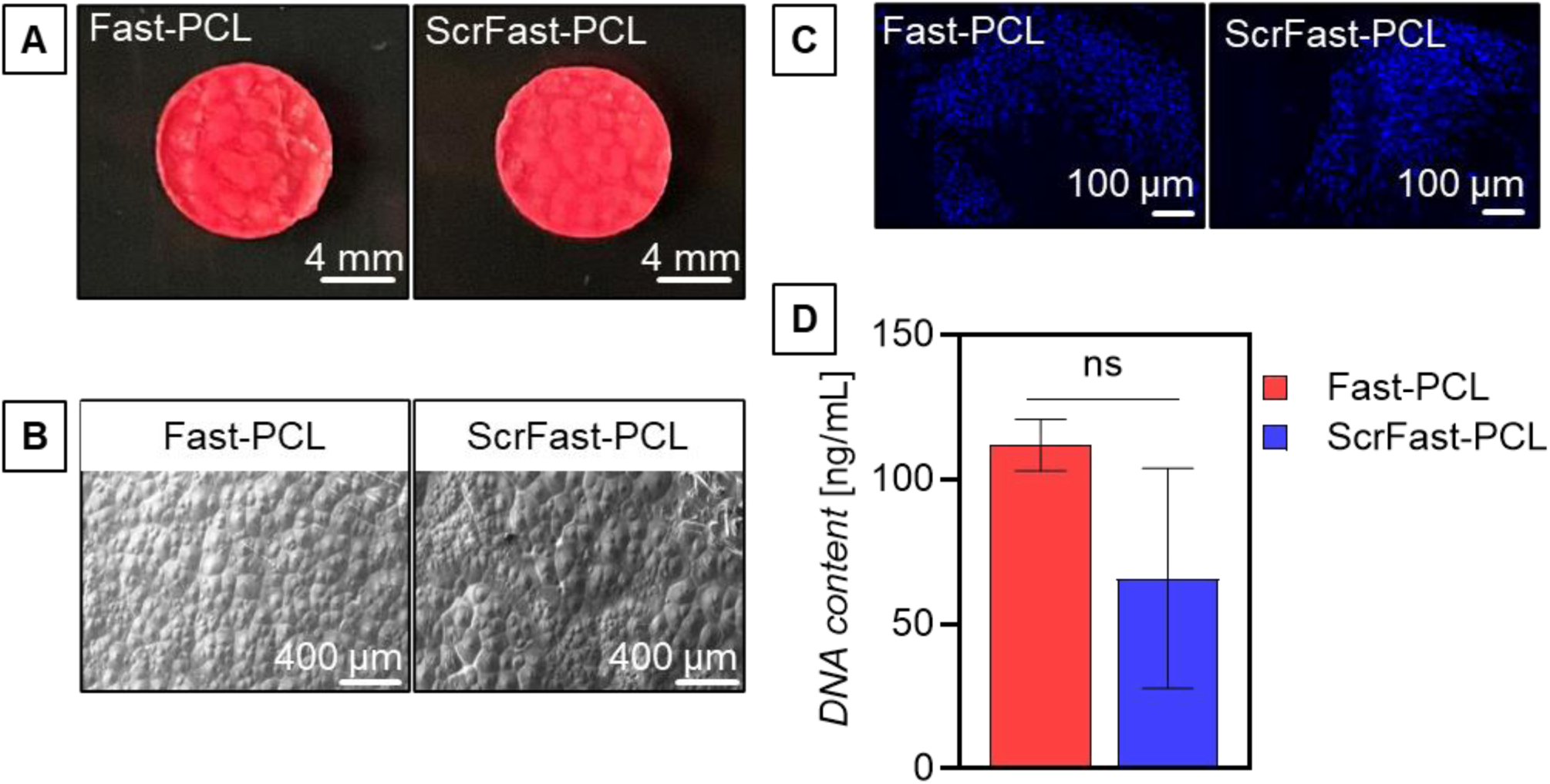
Representative (A) macroscopic and (B) scanning electron microscopy (SEM) images of solvent-cast scaffolds. (C) Representative fluorescence microscopy images of cell nuclei on peptide-PCL scaffolds stained with Hoechst 33342 (blue). (D) DNA quantification of hMSCs seeded on Fast-PCL and ScrFast-PCL scaffolds at Day 4 (N=3 scaffolds per group).

### 2.2. Scaffold degradation with collagenase

The degradation behavior of Fast-PCL and ScrFast-PCL scaffolds was assessed under proteolytic conditions to validate the platform design. Scaffold degradation was quantified via fluorescence intensity of Cy3 released into the media, providing a sensitive and real-time readout of peptide cleavage within the polymer backbone. We incubated the scaffolds in 2 mg/mL collagenase and monitored degradation for 21 days. Collagenase belongs to the MMP family and is crucial for ECM remodeling, owing to its ability to breakdown collagen, a primary ECM protein.^[69]^ Fluorescence measurements of collected solutions revealed that Fast-PCL scaffolds degraded significantly faster than ScrFast-PCL scaffolds in the presence of collagenase **(Figure 6)**. After 21 days, Fast-PCL scaffolds showed a cumulative Cy3 release of 20.8±2.17 nmol, equivalent to 25.77±3.70% of the total Cy3 in the scaffold, while ScrFast*-*PCL released only 8.70±0.92 nmol Cy3 release or 11.31±1.01 % total Cy3. Both scaffolds in buffer alone exhibited low levels of degradation, likely caused by non-specific hydrolysis of PCL ester bonds.^[18]^ Interestingly, ScrFast-PCL scaffolds degraded faster in collagenase compared to buffer alone. The presence of the PEG segments flanking the peptide sequences has been shown to increase hydrophilicity and protease accessibility.^[58–60]^ The ScrFast peptide showed slight degradation after 24 hours in the presence of multiple cell types (Figure 3), indicating some protease susceptibility that would be enhanced by the adjacent PEG spacers. We also observed Fast-PCL scaffolds in buffer degraded at the same rate as ScrFast-PCL scaffolds in collagenase but significantly faster than ScrFast-PCL scaffolds in buffer at all time points. Together, these results suggested that changes in amino acid sequence impact scaffold susceptibility to cleavage.

**Figure 6.**
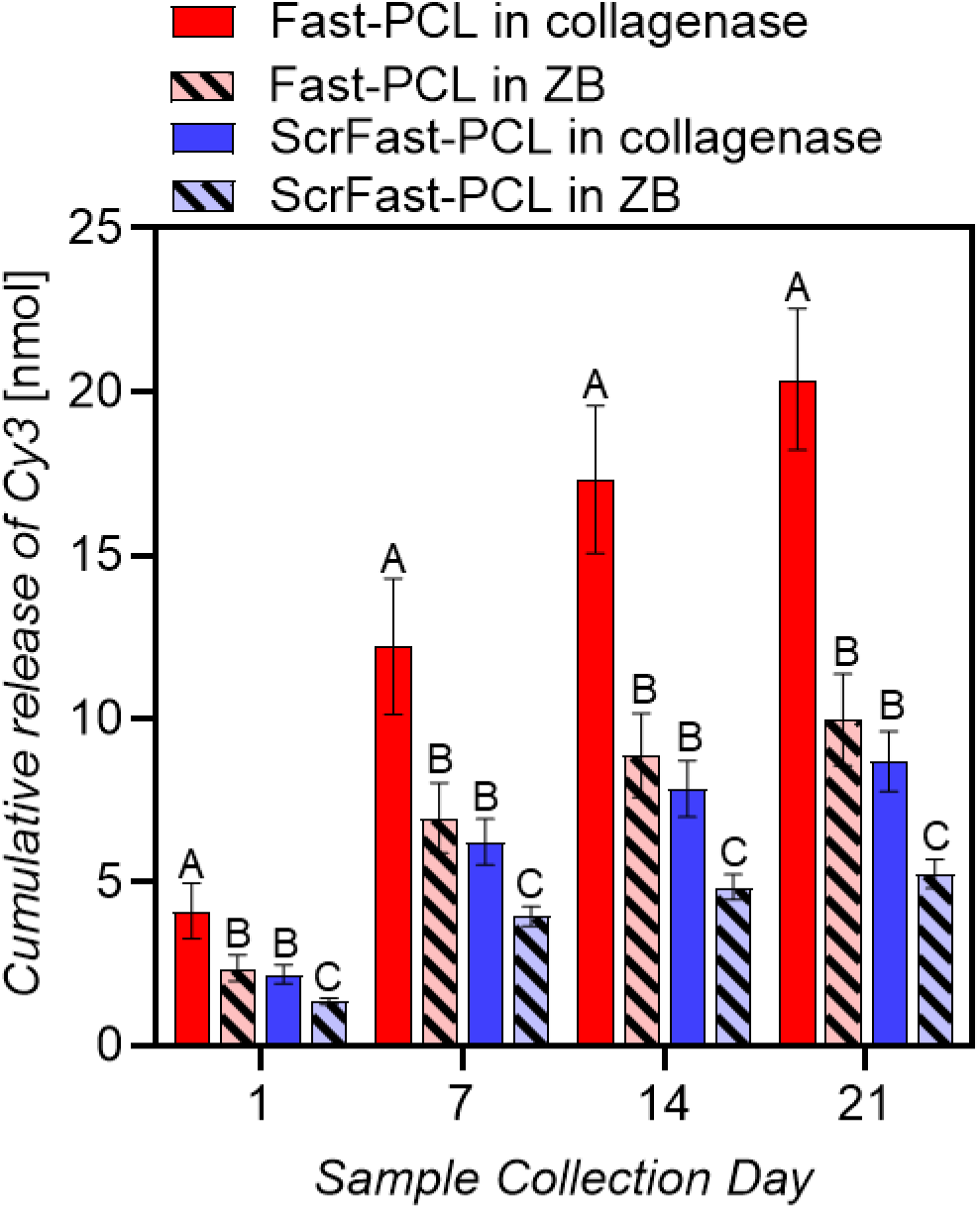
Collagenase-driven degradation of peptide-PCL scaffolds. Fluorescence analysis of collected media showed that scaffolds with a fast-degrading peptide sequence degraded significantly faster than scaffolds with a scrambled sequence of the same amino acids. Different letters within a timepoint indicate statistically significant differences (N=6 scaffolds per group, p < 0.05).

These findings verified that incorporating a protease-sensitive peptide into the PCL conjugate backbone resulted in protease-mediated scaffold degradation. Proteases were able to access and cleave peptides within the peptide-PCL conjugate backbone. In addition, cleavage likely generated shorter PCL fragments with reduced crystallinity and increased water infiltration, potentially leading to accelerated bulk degradation.^[70]^ Notably, the data confirmed successful translation of peptide degradation behavior from the peptide screening method to a scaffold material, highlighting sequence-specific selectivity of protease-mediated cleavage.

### 2.3. Cell-mediated scaffold degradation

Fast-PCL and ScrFast-PCL scaffolds were cultured with hMSCs for 10 days. MSCs play a key role in TE because they can be easily isolated from patients for autologous transplantation and are capable of differentiating into nearly all connective tissue phenotypes, including bone, cartilage, skeletal muscles, dense fibrous tissues (i.e. tendons and ligaments), and adipose tissue.^[71]^ In addition, hMSCs express proteases, including MMPs and plasmin, that can proteolytically degrade the peptides that are incorporated into the peptide-PCL conjugate scaffold.^[72]^

Media samples were collected throughout the culture period to quantify Cy3 release using fluorescence. We observed a higher amount of Cy3 released from Fast-PCL scaffolds (30.31±3.18 nmol Cy3 release; 26.68±2.17 % Cy3 total) compared to ScrFast-PCL scaffolds (23.91±2.13 nmol Cy3 release; 18.97±1.24 % Cy3 total), indicating Fast-PCL scaffolds degraded significantly faster compared to ScrFast-PCL scaffolds **(Figure 7A)**. We quantified DNA content for both scaffold groups compared to a tissue culture plastic (TCP) control at Days 4 and 10 to confirm that the significant differences in scaffold degradation were not caused by differences in cell number **(Figure 7B)**. In contrast, incubating the scaffolds in growth media without cells showed no measurable differences in degradation between Fast-PCL and ScrFast-PCL scaffolds. These data indicated that passive hydrolysis or media components alone did not account for the observed differences (**Figure 7A**). The Fast peptide was shown to be selectively cleaved by hMSC-secreted proteases while the scrambled control provided a baseline for nonspecific breakdown.

**Figure 7.**
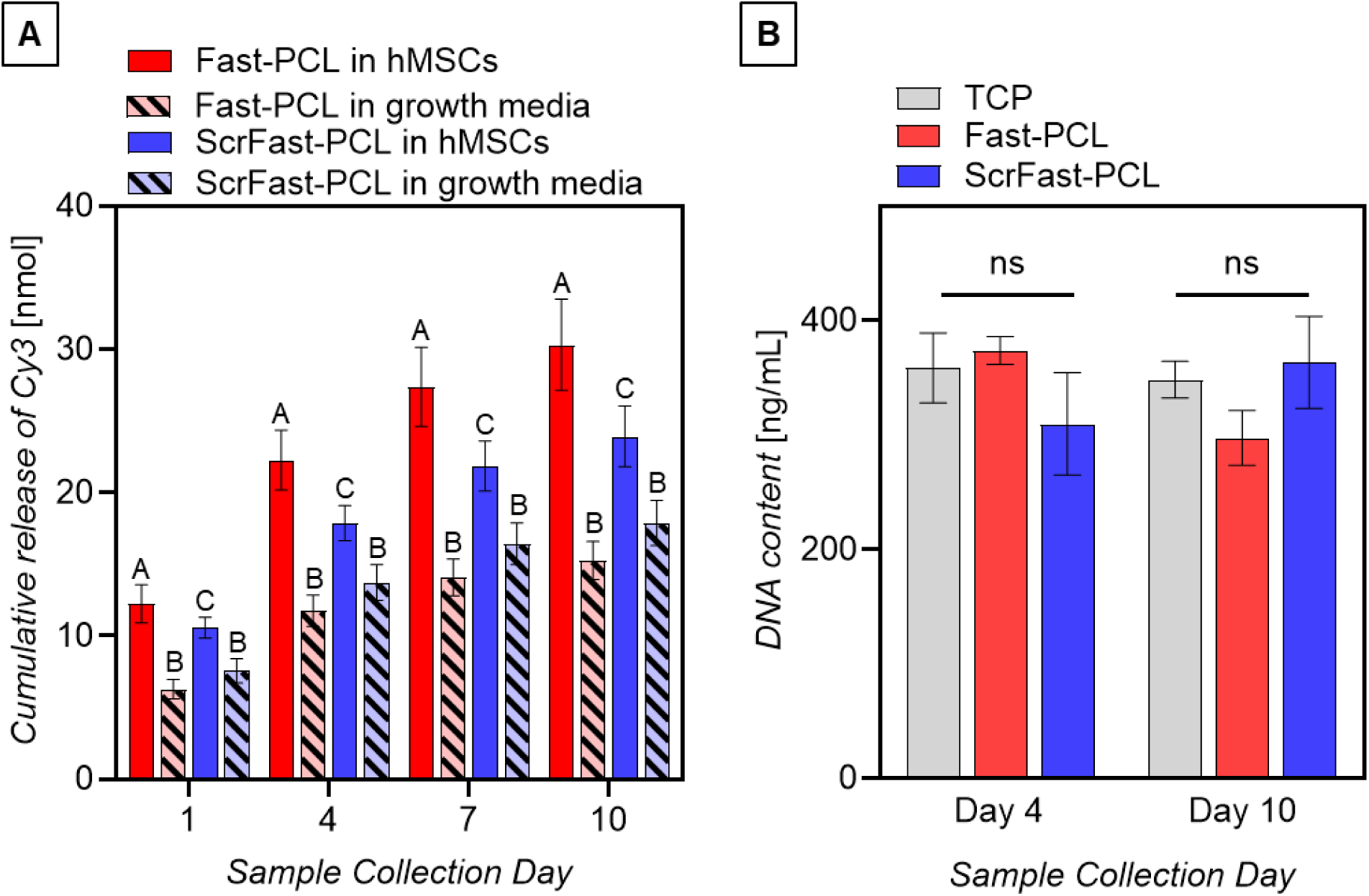
(A) hMSC-mediated degradation of peptide-PCL scaffolds. Quantification of Cy3 fluorescence in collected cell culture media showed that scaffolds with the Fast sequence degraded significantly faster than scaffolds with a scrambled sequence (ScrFast) containing the same amino acids. Different letters within a timepoint indicate statistically significant differences (N=5-6 scaffolds per group, p < 0.05). (B) DNA quantification of hMSCs cultured with Fast-PCL and ScrFast-PCL scaffolds at Days 4 and 10 showed no significant differences in cell number between scaffolds and the tissue culture plastic (TCP) control (N=3 scaffolds per group).

This study demonstrated that our protease-cleavable peptide-PCL conjugates underwent precise, cell-directed remodeling, establishing a new approach for engineering dynamic, resorbable solid scaffolds. We presented, to our knowledge, the first quantitative demonstration of cell-mediated, sequence-specific degradation in solid polymeric scaffolds by embedding a cleavage motif directly into the polymer backbone. This approach enabled scaffold degradation to be coupled to protease activity from surrounding cells. The data validated our proteomics-guided peptide discovery pipeline as a powerful strategy for designing biomaterials with sequence-specific degradation and position this platform for patient-tailored regenerative therapies.

## 3. Conclusion

We developed a versatile platform to fabricate solid polymeric scaffolds susceptible to cell-mediated degradation guided by protease activity. Using a proteomics-based approach, we identified a fast-degrading peptide sequence (Fast) cleaved by multiple cell types and designed a scrambled control sequence (ScrFast) resistant to proteolysis. Both peptides were integrated into the polycaprolactone (PCL) backbone to form linear peptide–PCL conjugates. Incorporating a Cy3 fluorophore into the conjugate enabled real-time, quantitative tracking of scaffold breakdown. In addition, PEG segments flanking the peptides ensured protease accessibility within the hydrophobic polymer scaffold. The Fast-PCL scaffolds degraded significantly faster than ScrFast scaffolds in response to collagenase and hMSCs, confirming that scaffold remodeling was both sequence-specific and protease-driven. These results validated that the peptides remained accessible within the polymer backbone and that scaffold degradation can be precisely tuned through peptide sequence design, establishing a modular framework for tailoring these biologically responsive materials.

This platform overcomes key limitations with protease-sensitive hydrogel systems, such as rapid degradation and limited mechanical properties, while retaining the processability and mechanical robustness of thermoplastic materials. The thiol-Michael addition conjugation strategy used to synthesize the peptide–PCL conjugates can be easily adapted for peptide sequences that target specific tissues and cell types. The conjugates are also compatible with other fabrication techniques, such as 3D printing, to generate patient-specific, free-standing 3D-printed constructs.^[11,68,73,74]^ Such scaffolds are particularly suited for orthopaedic and craniofacial applications where synchronizing implant degradation and bone tissue regeneration is critical for long-term success. Combining proteomics-guided peptide discovery with tunable conjugate chemistry establishes a foundation for next-generation biomaterials with biologically regulated remodeling to advance functional regenerative strategies.

## 4. Experimental Section

### 4.1 Materials

*Peptide synthesis.* Materials for peptide synthesis included fluorenylmethyloxycarbonyl chloride (Fmoc)-protected amino acids from AAPPTec and Fmoc-L-Lys(Boc)-OH sourced from CEM Corporation. Fmoc-Rink-amide 4-methylbenzhydryalmine (MBHA) resin and O-benzotriazole-N,N,N’,N’-tetramethyluronium hexafluoro-phosphate (HBTU) were also obtained from AAPPTec. Piperidine was acquired from BeanTown Chemical while N, N’-diisopropylethylamine (DIEA), trifluoroacetic acid (TFA), and triisopropylsilane (TIS) came from Sigma-Aldrich. Other solvents and reagents included diethyl ether (DEE), N,N-dimethylformamide (DMF), dichloromethane (DCM), and acetonitrile (ACN) from VWR; diisopropylcarbodiimide (DIC) from TCI America; ethyl 2-cyano-2(hydroxyamino)acetate (Oxyma) from CEM; and dithiothreitol (DTT) from Gold Biotechnology Inc. The ninhydrin test kit was purchased from Anaspec.

*Peptide-PCL conjugation.* Materials for conjugation included poly(caprolactone) (25 kDa) from Sigma-Aldrich and amine-poly(ethylene glycol)-maleimide (5 kDa) from Advanced BioChemicals (Lawrenceville, GA, USA). Anhydrous N-methyl pyrrolidone (NMP) was obtained from Alfa Aesar (Ward Hill, MA, USA), while dimethyl sulfoxide (DMSO) was purchased from VWR. Deuterated solvents for NMR analysis included DMSO-d_6_ and DCM-d_2_ from Sigma-Aldrich and DMF-d_7_ from Thermo Scientific Chemicals.

*Scaffold fabrication and sterilization.* Scaffold fabrication utilized 1,1,1,3,3,3-hexafluoro-2-propanol (HFIP) sourced from Matrix Scientific, ethanol from VWR, and phosphate buffer saline (PBS) tablets from Enzo Life Sciences. Additional materials included bovine serum albumin (BSA) from Sigma-Aldrich and Sylgard™ 184 silicone encapsulant from Electron Microscopy Sciences.

*Degradation with collagenase.* Collagenase Type I (Clostridium histolyticum) from Thermo Scientific Chemicals and Invitrogen™ Novex™ Zymogram Developing Buffer (10X) were purchased from Fisher Scientific.

*Cell culture, staining and DNA quantification.* THP-1 monocytes were obtained from ATCC. Human mesenchymal stromal cells (hMSCs) were obtained from RoosterBio, Inc. Fibroblasts from a 27-year-old Caucasian female were obtained from Promocell. Cell culture media and supplements included Dulbecco’s Modified Eagle’s Medium (DMEM; high glucose, GlutaMAX Supplement, pyruvate; Gibco™), fetal bovine serum (FBS; GeminiBio), antibiotic-antimycotic solution containing penicillin, streptomycin, and amphotericin B (anti/anti); Corning™), and RPMI Medium (Cytiva).

Other materials included L-ascorbic acid (Macron Fine Chemicals); interferon-γ (IFN-γ), macrophage colony-stimulating factor (M-CSF), interleukin-4 (IL-4), and interleukin-13 (IL-13) from Peprotech; and Gibco™ TrypLE™ Express Enzyme (1X), phenol red, and phorbol 12-myristate 13-acetate (PMA) from Fisher Scientific. L-glutamine was purchased from Gibco™ and lipopolysaccharide (LPS) was from Sigma. For nuclei staining, paraformaldehyde (PFA) was obtained from VWR and Hoechst 33342 from Sigma-Aldrich. DNA was quantified using an Invitrogen™ Quant-iT™ PicoGreen™ dsDNA assay kit purchased from Fisher Scientific and tris-EDTA buffer from Quality Biological.

### 4.2 Methods

*Peptide synthesis and purification.* Peptides (KGPQGIWGQK, KRVKRRLLMET (Fast), PPDKTSPEPA, LTRLMEKRRKV (ScrFast)) for the proteomic screening and sequence specificity study were synthesized on Fmoc-Rink-amide MBHA resin following standard Fmoc solid-phase peptide synthesis (SPPS) techniques using a CEM Liberty Blue automated microwave peptide synthesizer. The fast-degrading and scrambled fast-degrading peptides were modified with N- and C-terminal cysteine groups and a cyanine3 (Cy3) fluorophore to monitor scaffold degradation. These peptides were prepared with the following sequences: CK(Cy3)KRVKRRLLMETC (Fast) and CK(Cy3)LTRLMEKRRKVC (ScrFast). All amino acids except for the final two were added using a CEM Liberty Blue automated microwave peptide synthesizer. The cysteine and lysine (protected by a methyltrityl (Mtt) group) at the N-terminus were coupled manually. The Mtt group was removed using 2% (v/v) TFA and 5% (v/v) TIS in DCM. A water soluble Cy3 was coupled to the epsilon amine in the lysine side chain.

Manual SPSS was performed using a 100 mL peptide synthesis vessel and a wrist action shaker. Fmoc protecting groups were removed with 20% (v/v) piperidine in DMF and the resin was subsequently washed with DMF and DCM. Each amino acid coupling solution was prepared by dissolving 3.95 equivalents of HBTU with 4 molar equivalents of the Fmoc-protected amino acids in DMF. DIEA was added at 6 molar equivalents to the amino acid solution before adding to the resin. The reaction was left for at least 3 hours then thoroughly washed with DMF and DCM. Ninhydrin tests were conducted to verify successful Fmoc deprotection and amino acid coupling.

Automated SPSS was performed on an automated microwave peptide synthesizer following standard methods developed by CEM. Similar to manual SPSS, a 20% (v/v) piperidine in DMF was used for each deprotection step. Amino acid solutions (4-fold excess to resin) were prepared by dissolving in DMF. DIC and Oxyma, each at 10 molar equivalents, were used to activate the coupling reactions.

Peptides were cleaved from the resin using 95 %(v/v) TFA, 2.5 %(v/v) ultrapure water, 2.5 %(v/v) TIS, and 2.5 % (w/v) DTT. TFA was removed by rotary evaporation. Peptides were recovered by precipitation in cold DEE. The crude peptides were purified using reversed-phase high performance liquid chromatography (HPLC; Agilent 218 Prep HPLC, Agilent Technologies, Santa Clara, CA, USA). Peptides were dissolved at 10 mg/mL in 95 %(v/v) ultrapure water, 4.9% (v/v) ACN and 0.1% (v/v) TFA, sonicated and filtered through a 45 µm PTFE syringe filter to remove particulates. Samples were purified through an Agilent 5 Prep-C18 column (150 mm x 21.2 mm, 5 μm pore size, 100 Å particle size) using a mobile phase composed of ACN with 0.1% TFA and ultrapure water with 0.1% TFA. Fractions containing the desired peptide were pooled and subjected to rotary evaporation to remove ACN and TFA before lyophilization. The mass of purified peptides was confirmed by matrix-assisted laser desorption/ionization time of flight mass spectrometry (MALDI TOF-MS; Shimadzu Benchtop MALDI-ToF 8020).

*Peptide degradation studies with multiple cell types.* To generate macrophages, THP-1 cells were placed into a 48 well plate with a seeding density of 500,000 cells per well using 500 µL of RPMI containing 10% FBS and 1% anti-anti with PMA at a concentration of 100 ng/mL for two days to generate M0 macrophages. To induce M1 macrophage phenotypes, the media was changed after Day 2 to RPMI containing interferon-γ (IFN-γ), 20 ng/mL macrophage colony-stimulating factor (M-CSF), and 100 ng/mL lipopolysaccharide (LPS) and allowed to polarize for three days (Day 5 of culture). For M2 macrophages, the media was changed after Day 2 to RPMI containing 20 ng/mL M-CSF, 40 ng/mL of interleukin-4 (IL-4), and 20 ng/mL of interluekin-13 (IL-13). During the degradation studies, the media was changed to macrophage serum-free media with L-glutamine.

Human MSCs from a 19-year-old Eritrean/East African male donor and THP-1 derived macrophages were seeded in 48-well plates (36,000 per well) while fibroblasts were seeded in 96-well plates (10,000 per well). The cells were cultured in 500 µL medium per well before replacing with fresh media after 24 hours. Each peptide was added at 50 µM to the media. A non-proteolytically degradable NH_2_-βFβAβAβAβAβAβA-amide peptide (βFβA_6_) (βF is β-phenylalanine and βA is β-alanine) was added at 50 µM to serve as an internal standard. Conditioned media (40 µL) was sampled at 0 and 24 hours. Samples were stored in a −80 °C freezer until analysis to prevent further proteolytic degradation. Samples were thawed and analyzed by liquid chromatography-mass spectrometry (LC-MS). Acetic acid (4 µL) was added to all collected media in the LC-MS plates. Peptides degradation ratios were measured by LC-MS (Thermo Fisher Vanquish UPLC, LTQ-XL mass spectrometer) and analyzed by Xcalibur. The relative concentrations of peptide were calculated by the ratio of peak area of peptide and peak area of the internal standard βFβA_6_.

*Synthesis of peptide-PCL conjugates.* Polycaprolactone (25 kDa; PCL) was modified with amine-poly(ethylene glycol)-maleimide (5 kDa; amine-PEG-mal) following an amide coupling process. PCL (125 mg/mL) was dissolved in anhydrous NMP. HBTU was added as an activator at a mol ratio of 4:3.95 PCL:HBTU. DIEA was added to the PCL/activator solution at a molar ratio of 4:6 PCL:DIEA to activate the carboxyl end of PCL. Amine-PEG-mal dissolved in 50:50 NMP:DMSO at 2 molar equivalents to PCL was added dropwise to the activated PCL solution while stirring at 350 rpm. The resulting solution was purged with N_2_ for 15 minutes and left to stir overnight while protected from light. Maleimide-terminated PCL (PCL-PEG-mal) was recovered by trituration with room temperature DEE.

Peptide-PCL (Fast-PCL and ScrFast-PCL) conjugates were synthesized via maleimide-thiol conjugation between PCL-PEG-mal and the thiol-terminated peptides. PCL-PEG-mal was dissolved in anhydrous NMP at 2 molar equivalents to the peptide. The peptide was separately dissolved in anhydrous NMP then added dropwise to the PCL-PEG-mal solution while stirring at 500 rpm. The resulting solution was purged with N_2_ for 15 minutes and continued to react overnight while protected from light. Fast-PCL and ScrFast-PCL conjugates were obtained by trituration with cold DEE. RGDS-PCL conjugate was also prepared using established methods previously described.^[68]^ Each synthesis was confirmed by proton nuclear magnetic resonance (^1^H NMR) spectroscopy. NMR samples were prepared by dissolving in either DCM-d_2_, DMSO-d_6_ or DMF-d_7_ and analyzed using a 500 MHz NMR spectrometer (Bruker).

*Scaffold fabrication.* Inks were prepared by mixing Fast-PCL or ScrFast-PCL (195 mg/mL) and RGDS-PCL (5 mg/mL) conjugates in HFIP. Inks were shaken on a wrist-action shaker at room temperature for 48 hours then allowed to rest at room temperature for 24 hours. Inks were cast onto the top lid of a 60 mm glass petri dish and allowed to dry in the hood for approximately 45 minutes. To ensure flatness, the bottom lid was placed over the dried ink, taped, and left to dry overnight. Scaffolds were then punched into 10 mm disks using a biopsy punch. The disks were immersed in 75% ethanol for 30 minutes, rinsed three times with sterile ultrapure water, soaked in 0.1% BSA for at least four hours, and subsequently rinsed three times with sterile PBS. The scaffolds were left to dry inside a biosafety hood before use.

*SEM characterization.* Scaffold surfaces were characterized using a scanning electron microscope (SEM; Axia ChemiSEM, ThermoFisher Scientific). Samples were mounted on 12-mm aluminum stubs using carbon tape then coated with iridium using a sputter coater (Electron Microscopy Sciences). Sample images were obtained using a secondary electron detector with an accelerating voltage of 5 kV.

*Cell adhesion assay.* Human MSCs from a 25-year-old Caucasian male donor were used to assess cell adhesion on Fast-PCL and ScrFast-PCL scaffolds. These cells were cultured in DMEM-GlutaMAX™ supplemented with 10% (v/v) FBS, 1% (v/v) anti/anti solution, and 0.1% L-ascorbic acid and maintained at 37 °C in 5% CO_2_. Cells were expanded to 80-90% confluency and subsequently harvested using Gibco™ TrypLE™ Express Enzyme (1X), phenol red. Passages 3-4 were selected for cell seeding. Cells (50,000 cells/mL; 100 µL) were seeded on top of the pinned scaffolds and incubated at 37 °C and 5% CO_2_ to allow for cell adhesion. After one hour, growth media was added in each well to reach a final volume of 1 mL. Samples were harvested and analyzed for fluorescence imaging on Day 4. Culture media was removed from the wells and samples were rinsed twice with sterile PBS. Samples were fixed in 4% (w/v) PFA in PBS for 1 hour at 4°C then washed two times in PBS. All samples were stored in PBS at 4°C until stained.

Prior to staining, cells were permeabilized using 0.1% (v/v) Triton X-100 in PBS for 15 minutes. Hoechst 33342 (10 mg/mL in ultrapure water) diluted 1:5000 in ultrapure water was added for nuclear staining. Samples were washed twice with PBS and stored in PBS at 4°C protected from light until imaging. Fluorescence images were acquired using a Keyence BZ-X810 microscope.

*In vitro degradation with collagenase.* Scaffolds were pinned to silicone elastomer-coated 24-well plates using 0.1 mm dissection pins. The degradation behavior of the scaffolds in the presence of collagenase was assessed by immersing pinned scaffolds in 2 mg/mL collagenase in Zymogram-developing buffer (ZB). Solutions (1 mL) were collected and changed after the first day of incubation then after every 3-4 days. Samples were placed in an incubator (37 °C, 5% CO_2_) for up to 21 days. Scaffolds were also exposed to ZB as a baseline control.

*In vitro degradation with hMSCs.* Human MSCs were cultured as described above to assess cell-mediated degradation of Fast-PCL and ScrFast-PCL scaffolds *in vitro*. Cells were seeded at passage 3-4 into 24-well plates at 50,000 cells/well in 100 µL and incubated at 37 °C with 5% CO_2_ for one hour to allow for cell adhesion. Media was added to each well to a final volume of 1 mL per well. Culture media was replaced after the first day and then every three days to monitor degradation for up to 10 days. Scaffolds were also immersed in growth media alone as a baseline control.

*DNA quantification.* DNA content was analyzed on Days 4 and 10. After collecting media, samples were carefully rinsed twice with sterile PBS. Subsequently, 0.2% Triton X-100 (0.5 mL) was added to each well, and cells were scraped and thoroughly mixed using a pipette. The resulting solutions were transferred to Eppendorf tubes and stored at −80 °C until further analysis. DNA quantification was carried out using the PicoGreen assay, with fluorescence measurements taken at excitation and emission wavelengths of 485 nm and 520 nm, respectively, using an Infinite M Nano Tecan microplate reader. DNA ladder (0-1000 ng/mL) was prepared using calf thymus DNA in 1X Tris-EDTA buffer.

*Fluorescence analysis.* The degradation of Fast-PCL and ScrFast-PCL scaffolds was assessed by quantifying the Cy3 released into the solution. Fluorescence intensity was measured with excitation and emission wavelengths set at 550 nm and 580 nm, respectively, using an Infinite M Nano Tecan microplate reader (Tecan Austria GmbH). A standard curve (0 – 2 nmol Cy3) was created using Cy3 dissolved in ZB, collagenase in ZB, or hMSC growth media to calculate the Cy3 released by the scaffolds in different conditions.

*Statistical analysis.* Statistical analysis was performed using SPSS (IBM, version 31.0.0.0). Independent samples t-tests were used to compare the degradation behavior of Fast and ScrFast peptides across different cell types (N = 3) and to assess cell attachment on Fast-PCL and ScrFast-PCL scaffolds (N = 3). Statistical significance for these experiments was defined as *p < 0.05, **p < 0.01, and ***p < 0.001. For comparisons involving more than two groups, one-way analysis of variance (ANOVA) was applied followed by Tukey’s HSD or Dunnett’s T3 post-hoc tests as appropriate. This analysis was used to compare relative fluorescence intensity of solutions collected at different time points (N = 5–6 scaffolds per group) and DNA content per timepoint (N = 3). Groups not sharing the same letter were considered significantly different (p < 0.05).

## Supporting Information

Supporting Information is available from the Wiley Online Library or from the author.

## Acknowledgements

The authors acknowledge Lehigh’s Electron Microscopy and Nanofabrication Facility, Institute for Functional Materials and Devices (I-FMD), Health Research Hub (HRH) and Health, Science, and Technology (HST) facilities, and shared instrumentation in the Department of Chemistry. This work was generously supported by a Collaborative Research Grant from Lehigh University awarded to LWC and ETP and partially supported by the National Science Foundation (NSF) through a Faculty Early Career Development (CAREER) award (DMR 1944914 to LWC) and grants from the Musculoskeletal Transplant Foundation to WLG and the National Institutes of Health (NIH) National Institute of Dental and Craniofacial Research (NIDCR) (1R01DE027957 to WLG). The Bruker Neo 500 MHz NMR was acquired through NSF-MRI-1725883 with additional support from Lehigh University.

## Author Contributions

K.V.G.S. did the study design, performed experiments, data analysis and interpretation, wrote the original draft of the manuscript, and reviewed and edited the final manuscript. N.K.H. conceptualized the project, performed preliminary experiments, and contributed to manuscript editing. S.S. conceptualized the project, contributed to manuscript editing, and provided critical review comments. K.B.S. conceptualized the project and performed preliminary experiments. Y.W. performed experiments, data analysis, and interpretation. E.T.P. conceptualized the project, performed experiments, data analysis, and interpretation, contributed to manuscript writing, and provided critical review comments. W.L.G. conceptualized the project, helped with data analysis and interpretation, contributed to manuscript editing, and provided critical review comments. L.W.C. conceptualized and supervised the project, secured funding, performed data analysis and interpretation, contributed to manuscript writing, and provided critical review comments. All authors reviewed and approved the final version of the manuscript.

## Data Availability Statement

The data that support the findings of this study are available from the corresponding author upon reasonable request.

## References

[1] F. J. O’Brien, Mater. Today 2011, 14, 88.

[2] F. Zhang, M. W. King, Adv. Healthcare Mater. 2020, 9, 1901358.

[3] A. E. Eldeeb, S. Salah, N. A. Elkasabgy, AAPS PharmSciTech 2022, 23, 267.

[4] A. Kirillova, T. R. Yeazel, D. Asheghali, S. R. Petersen, S. Dort, K. Gall, M. L. Becker, Chem. Rev. 2021, 121, 11238.

[5] A. K. Gaharwar, I. Singh, A. Khademhosseini, Nat. Rev. Mater. 2020, 5, 686.

[6] M. Bartnikowski, T. R. Dargaville, S. Ivanovski, D. W. Hutmacher, Prog. Polym. Sci. 2019, 96, 1.

[7] Y. You, B. M. Min, S. J. Lee, T. S. Lee, W. H. Park, J. Appl. Polym. Sci. 2005, 95, 193.

[8] Z. Terzopoulou, A. Zamboulis, I. Koumentakou, G. Michailidou, M. J. Noordam, D. N. Bikiaris, Biomacromolecules 2022, 23, 1841.

[9] L. L. Osorno, A. N. Brandley, D. E. Maldonado, A. Yiantsos, R. J. Mosley, M. E. Byrne, Nanomaterials 2021, 11, 278.

[10] F. Rey, B. Barzaghini, A. Nardini, M. Bordoni, G. V. Zuccotti, C. Cereda, M. T. Raimondi, S. Carelli, Cells 2020, 9, 1636.

[11] J. W. Tolbert, D. E. Hammerstone, N. Yuchimiuk, J. E. Seppala, L. W. Chow, Macromol. Mater. Eng. 2021, 306, 202100442.

[12] M. Gharibshahian, M. Salehi, N. Beheshtizadeh, M. Kamalabadi-Farahani, A. Atashi, M. S. Nourbakhsh, M. Alizadeh, Front. Bioeng. Biotechnol. 2023, 11, 1168504.

[13] F. Sun, X. Sun, H. Wang, C. Li, Y. Zhao, J. Tian, Y. Lin, Int. J. Mol. Sci. 2022, 23, 5831.

[14] E. Salamanca, C. S. Choy, L. M. Aung, T. C. Tsao, P. H. Wang, W. A. Lin, Y. F. Wu, W. J. Chang, Polymers 2023, 15, 2619.

[15] N. Sagar, B. Chakravarti, S. S. Maurya, A. Nigam, P. Malakar, R. Kashyap, Front. Bioeng. Biotechnol. 2024, 12, 1385365.

[16] S. Singh, E. L. Nyberg, A. N. O’Sullivan, A. Farris, A. N. Rindone, N. Zhang, E. C. Whitehead, Y. Zhou, E. Mihaly, C. C. Achebe, W. Zbijewski, W. Grundy, D. Garlick, N. D. Jackson, T. Taguchi, C. Takawira, J. Lopez, M. J. Lopez, M. P. Grant, W. L. Grayson, Biomaterials 2022, 282, 121392.

[17] E. Nyberg, A. Farris, A. O’Sullivan, R. Rodriguez, W. Grayson, Tissue Eng. Part A 2019, 25, 1459.

[18] L. N. Woodard, M. A. Grunlan, ACS Macro Lett. 2018, 7, 976.

[19] T. Honma, T. Itagaki, M. Nakamura, S. Kamakura, I. Takahashi, S. Echigo, Y. Sasano, Oral Dis. 2008, 14, 457.

[20] S. Bose, M. Roy, A. Bandyopadhyay, Trends Biotechnol. 2012, 30, 546.

[21] H. Ramaraju, A. S. Verga, B. J. Steedley, A. P. Kowblansky, G. E. Green, S. J. Hollister, Biomaterials 2025, 321, 123257.

[22] M. J. Dewey, B. A. C. Harley, RSC Adv. 2021, 11, 17809.

[23] S. Abdulghani, G. R. Mitchell, Biomolecules 2019, 9, 750.

[24] S. Tajvar, A. Hadjizadeh, S. S. Samandari, Int. Biodeterior. Biodegrad. 2023, 180, 105599.

[25] H. Zhang, L. Zhou, W. Zhang, Tissue Eng. Part B Rev 2014, 20, 492.

[26] A. S. Sawhney, J. A. Hubbell, J. Biomed. Mater. Res. 1990, 24, 1397.

[27] M. Malin, M. Hiljanen-Vainio, T. Karjalainen, J. Seppälä, J. Appl. Polym. Sci. 1996, 59, 1289.

[28] S. A. Eming, P. Martin, M. Tomic-Canic, Sci. Transl. Med. 2014, 6, 265sr6.

[29] M. H. Yun, Int. J. Mol. Sci. 2015, 16, 25392.

[30] M. S. Ghiasi, J. Chen, A. Vaziri, E. K. Rodriguez, A. Nazarian, Bone Rep. 2017, 6, 87.

[31] V. Fischer, M. Kalbitz, F. Müller-Graf, F. Gebhard, A. Ignatius, A. Liedert, M. Haffner-Luntzer, Int. J. Mol. Sci. 2018, 19, 2070.

[32] A. Karpouzos, E. Diamantis, P. Farmaki, S. Savvanis, T. Troupis, J Osteoporos. 2017, 4218472.

[33] E. Gibon, F. Loi, L. A. Córdova, J. Pajarinen, T. Lin, L. Lu, A. Nabeshima, Z. Yao, S. B. Goodman, Regen. Eng. Transl. Med. 2016, 2, 98.

[34] I. Pountos, T. Georgouli, H. Bird, G. Kontakis, P. V. Giannoudis, Expert Opin. Drug Saf. 2011, 10, 935.

[35] J. Losner, K. Courtemanche, J. L. Whited, npj Regener. Med. 2021, 6, 1.

[36] M. Marenzana, T. R. Arnett, Bone Res 2013, 1, 203.

[37] A. Zengin, F. C. Teixeira, T. Feliciano, P. Habibovic, C. D. Mota, M. B. Baker, S. van Rijt, Biomater. Adv. 2023, 154, 213647.

[38] D. Gräfe, M. Gernhardt, J. Ren, E. Blasco, M. Wegener, M. A. Woodruff, C. Barner-Kowollik, Adv. Funct. Mater. 2021, 31, 2006998.

[39] Y. Li, M. D. Hoffman, D. S. W. Benoit, Biomaterials 2020, 268, 120535.

[40] A. Yahyouche, X. Zhidao, J. T. Czernuszka, A. J. P. Clover, Acta Biomater. 2011, 7, 278.

[41] J. L. West, J. A. Hubbell, Macromolecules 1999, 32, 241.

[42] T. H. Qazi, M. R. Blatchley, M. D. Davidson, F. M. Yavitt, M. E. Cooke, K. S. Anseth, J. A. Burdick, Cell Stem Cell 2022, 29, 678.

[43] P. Lu, K. Takai, V. M. Weaver, Z. Werb, Cold Spring Harbor Perspect. Biol. 2011, 3, a005058.

[44] M. B. Pimentel, F. T. P. Borges, F. Teymour, O. Y. Zaborina, J. C. Alverdy, K. Fang, S. H. Hong, A. Staneviciute, Y. J. He, G. Papavasiliou, J. Mater. Chem. B 2020, 8, 2454.

[45] Y. Rong, Z. Zhang, C. L. He, X. S. Chen, Sci. China Technol. Sci. 2021, 64, 1285.

[46] R. R. Katz, J. L. West, Adv. Biol. 2022, 6, 2200084.

[47] M. S. Mazzeo, T. Chai, M. Daviran, K. M. Schultz, ACS Appl. Bio. Mater. 2018, 2, 81.

[48] S. Sokic, G. Papavasiliou, Tissue Eng. Part A 2012, 18, 2477.

[49] B. V. Sridhar, J. L. Brock, J. S. Silver, J. L. Leight, M. A. Randolph, K. S. Anseth, Adv Healthcare Mater. 2015, 4, 702.

[50] J. Guo, H. Sun, W. Lei, Y. Tang, S. Hong, H. Yang, F. R. Tay, C. Huang, J. Dent. Res. 2019, 98, 564.

[51] X. Wei, S. Chen, T. Xie, H. Chen, X. Jin, J. Yang, S. Sahar, H. Huang, S. Zhu, N. Liu, C. Yu, P. Zhu, W. Wang, W. Zhang, Theranostics 2022, 12, 127.

[52] K. B. Fonseca, D. B. Gomes, K. Lee, S. G. Santos, A. Sousa, E. A. Silva, D. J. Mooney, P. L. Granja, C. C. Barrias, Biomacromolecules 2014, 15, 380.

[53] R. Chandrawati, Exp. Biol. Med. 2016, 241, 972.

[54] G. A. Foster, D. M. Headen, C. González-García, M. Salmerón-Sánchez, H. Shirwan, A. J. García, Biomaterials 2016, 113, 170.

[55] S. L. Fung, J. P. Cohen, E. T. Pashuck, C. E. Miles, J. W. Freeman, J. Kohn, J. Mater. Chem. B 2023, 11, 6621.

[56] G. Zorn, F. I. Simonovsky, B. D. Ratner, D. G. Castner, Adv. Healthcare Mater. 2021, 11, 2100894.

[57] K. S. Stakleff, F. Lin, L. A. Smith Callahan, M. B. Wade, A. Esterle, J. Miller, M. Graham, M. L. Becker, Acta Biomater. 2013, 9, 5132.

[58] J. Guan, W. R. Wagner, Biomacromolecules 2005, 6, 2833.

[59] H. L. Fu, Y. Hong, S. R. Little, W. R. Wagner, Biomacromolecules 2014, 15, 2924.

[60] H. Benhardt, N. Sears, T. Touchet, E. Cosgriff-Hernandez, Macromol. Biosci. 2011, 11, 1020.

[61] M. Ramezani, M. B. B. Monroe, J. Biomed. Mater. Res. A 2023, 111, 921.

[62] Y. Wu, S. J. Rozans, A. S. Moghaddam, E. T. Pashuck, Adv. Healthcare Mater. 2025, e01932.

[63] S. J. Rozans, Y. Wu, A. S. Moghaddam, E. T. Pashuck, J. Biomed. Mater. Res. A 2025, 113, e37864.

[64] J. Patterson, J. A. Hubbell, Biomaterials 2010, 31, 7836.

[65] C. E. Hoyle, A. B. Lowe, C. N. Bowman, Chem. Soc. Rev. 2010, 39, 1355.

[66] B. E. Turk, L. C. Cantley, Methods 2004, 32, 398.

[67] T. Klein, U. Eckhard, A. Dufour, N. Solis, C. M. Overall, Chem. Rev. 2018, 118, 1137.

[68] P. Camacho, H. Busari, K. B. Seims, P. Schwarzenberg, H. L. Dailey, L. W. Chow, Biomater. Sci. 2019, 7, 4237.

[69] Z. Z. Wang, K. Wang, L. F. Xu, C. Su, J. S. Gong, J. S. Shi, X. D. Ma, N. Xie, J. Y. Qian, BioDesign Res. 2024, 6, 0050.

[70] A. Leroux, T. Ngoc Nguyen, A. Rangel, I. Cacciapuoti, D. Duprez, D. G. Castner, V. Migonney, Biointerphases 2020, 15, 061006.

[71] N. W. Marion, J. J. Mao, Methods Enzymol. 2006, 420, 339.

[72] T. P. Lozito, W. M. Jackson, L. J. Nesti, R. S. Tuan, Matrix Biol. 2014, 34, 132.

[73] P. Camacho, A. Behre, M. Fainor, K. B. Seims, L. W. Chow, Biomater. Sci. 2021, 9, 6813.

[74] P. Camacho, M. Fainor, K. B. Seims, J. W. Tolbert, L. W. Chow, J. Biol. Methods 2021, 8, 1.

