## Supplementary Information for "Design and Synthesis of Peptide-Polyester Conjugates for Cell-Mediated Scaffold Degradation"

**Figure S1.** Purified CK(Cy3)KRVKRRLLMETC. Representative (A) $^{1}H$ NMR spectrum (500 MHz, DMSO-$d_6$) and corresponding chemical structures: $\delta$ = 8.36 (t, 1H, vinyl H, a), 7.65 (d, 2H, Ar-H, b), 7.53 (d, 2H, Ar-H, c), 7.45 (m, 2H, Ar-H, d), 7.31 (t, 2H, Ar-H, e), and 0.81 (m,18H, -$CH_3$, 1), and (B) MALDI-ToF MS spectrum (MW 2311 Da), and (C) HPLC profile.

A

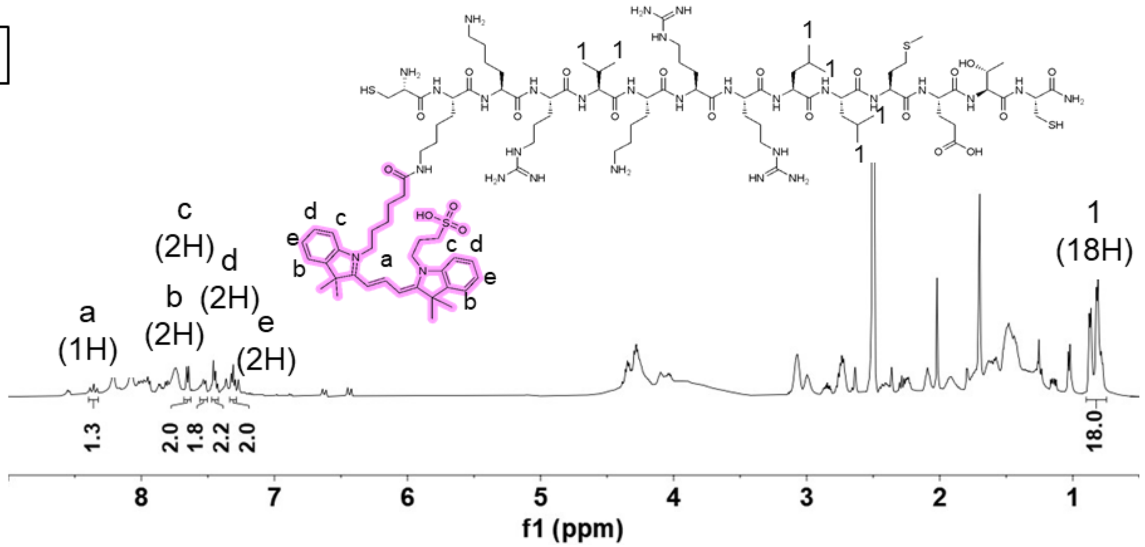


B

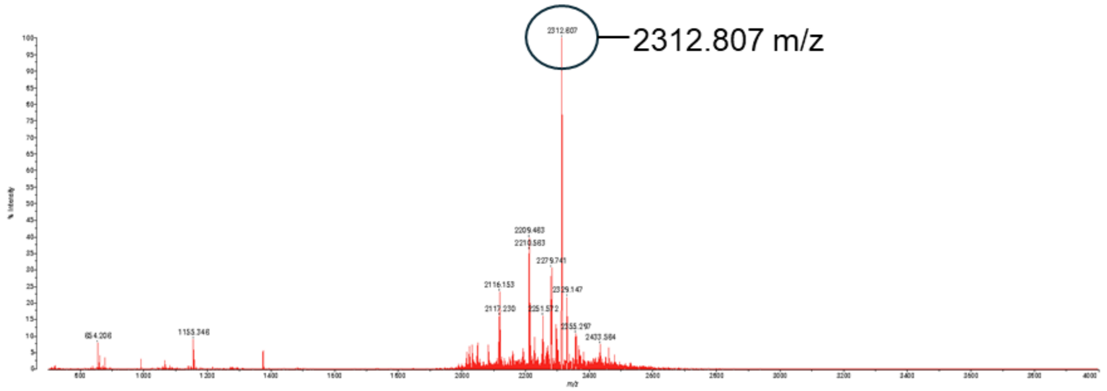


C

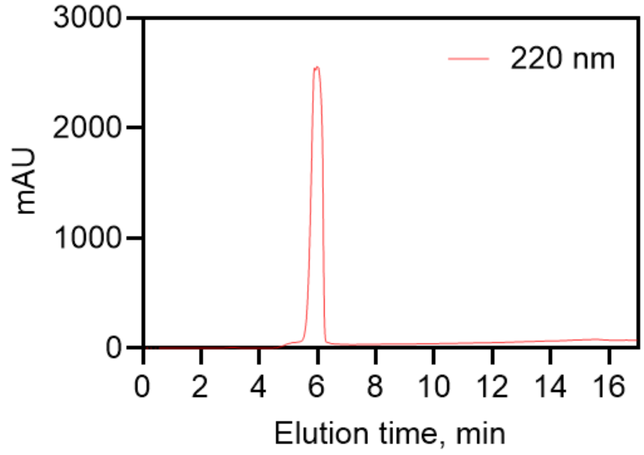

**Figure S2.** Purified CK(Cy3)LTRLMEKRRKVC. Representative (A) $^{1}$H NMR spectrum (500 MHz, DMSO-d$_6$) and corresponding chemical structures: δ = 8.35 (t, 1H, vinyl H, a), 7.64 (d, 2H, Ar-H, b), 7.53 (d, 2H, Ar-H, c), 7.44 (m, 2H, Ar-H, d), 7.30 (t, 2H, Ar-H, e), and 0.84 (m,18H, -CH$_3$, 1), and (B) MALDI-ToF MS spectrum (MW 2311 Da), and (C) HPLC profile.

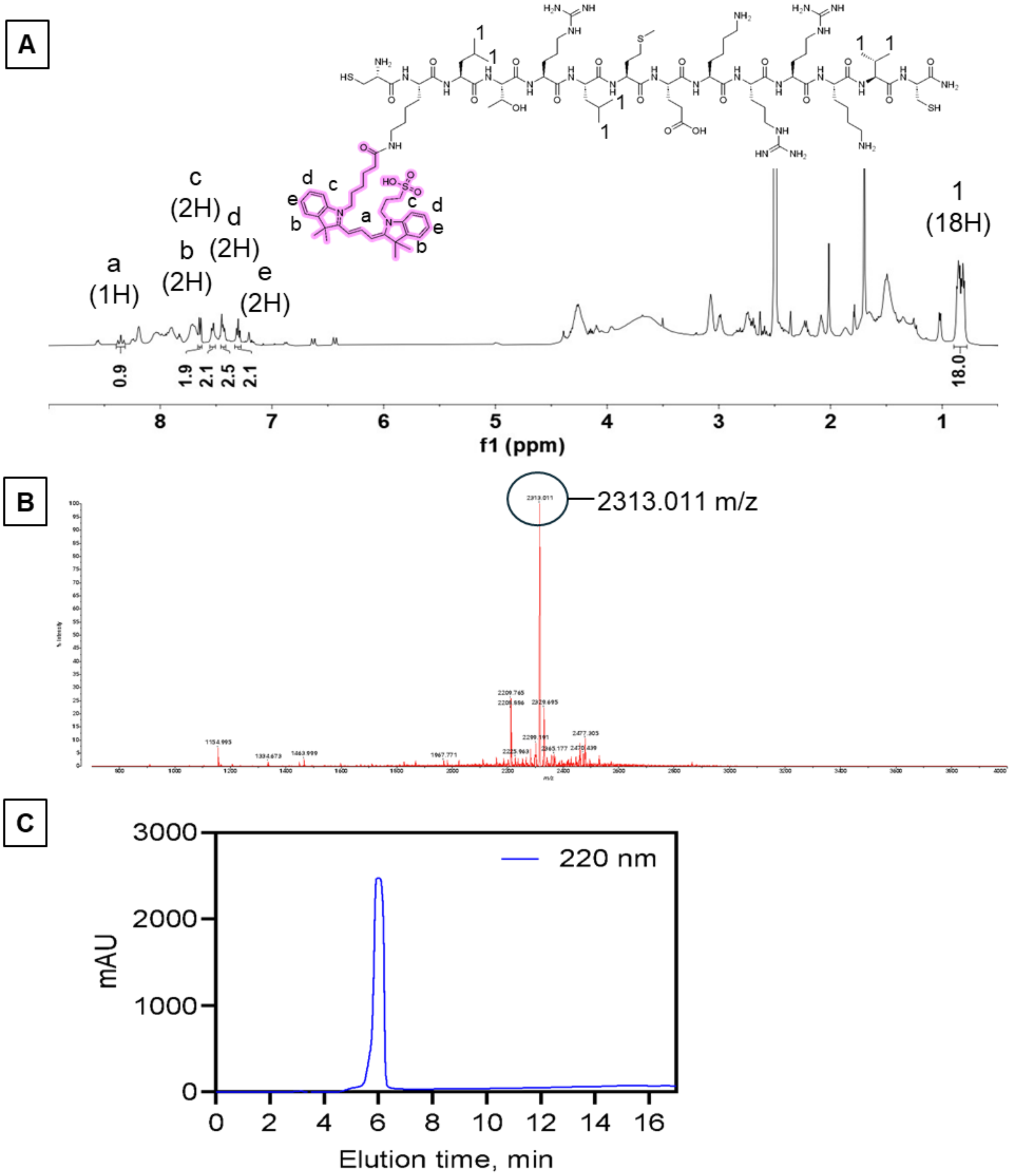

**Figure S3.** Peptide-PCL conjugation reaction scheme. Peptide was either a modified fast-degrading peptide sequence (Fast; CK(Cy3)KRVKRRL↓LMETC) or a scrambled version ScrFast; CK(Cy3)LTRLMEKRRKVC).

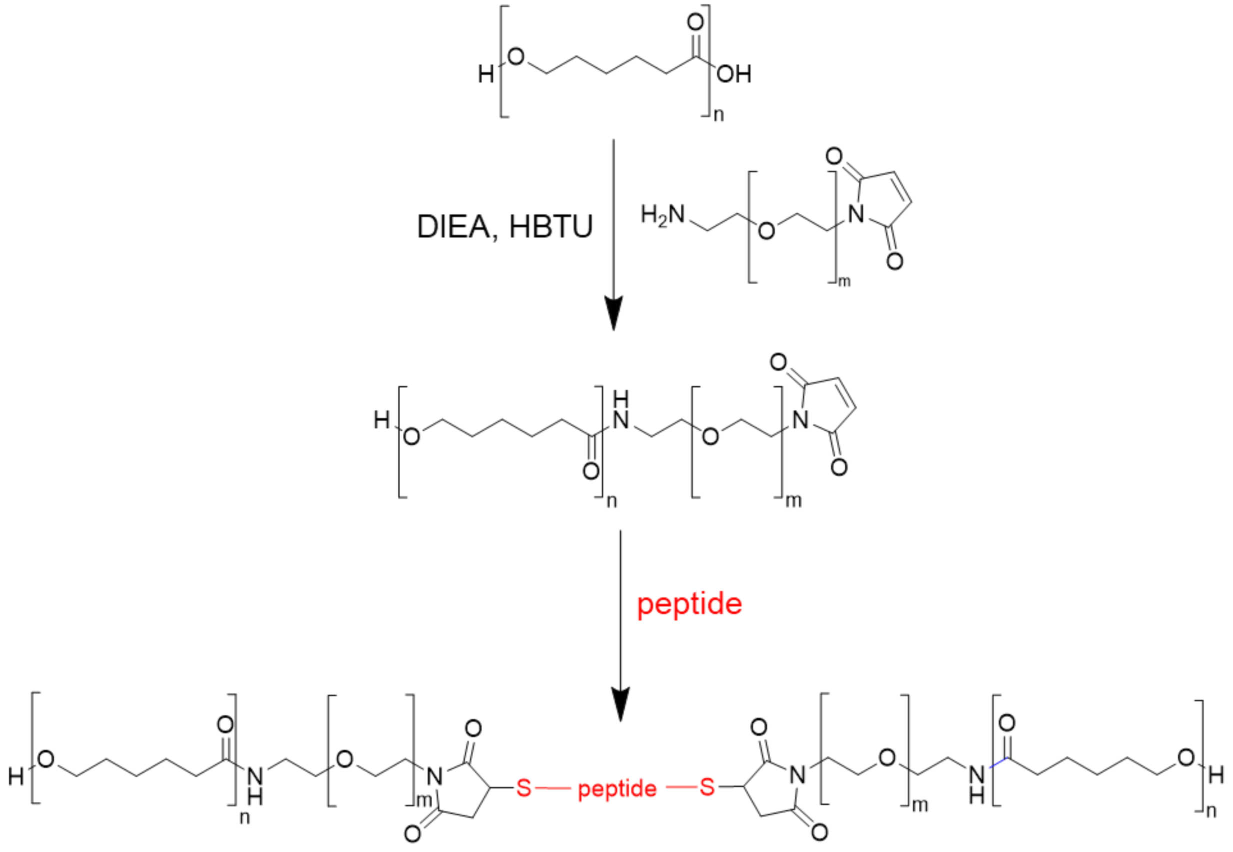

**Figure S4.** Representative $^{1}H$ NMR spectrum (500 MHz, DCM-$d_2$) and corresponding chemical structures of maleimide-functionalized PCL: δ = 6.69 (s, 2H, maleimide vinyl H, 1) and 4.03 (t, 456H, -O-$CH_2$-, a).

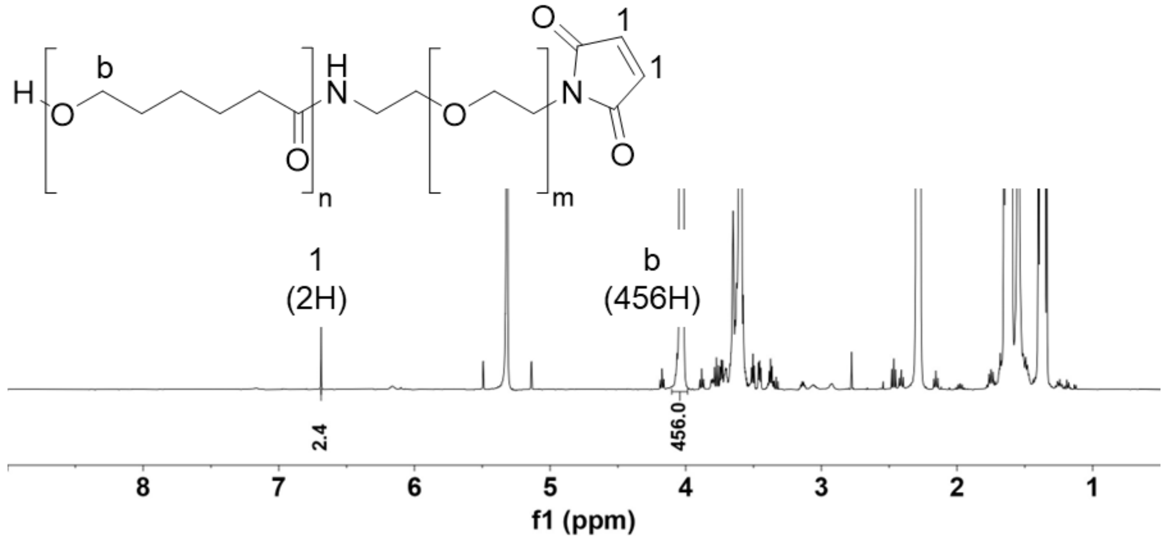

**Figure S5.** Purified CYGGGRGDS (RGDS). Representative (A) $^1$H NMR spectrum (500 MHz, DMSO-d$_6$) and corresponding chemical structures with chemical shift assignments: 6.65 (d, 2H, Ar-H, 1), (B) MALDI-ToF MS spectrum (MW 869 Da), and (C) HPLC profile.

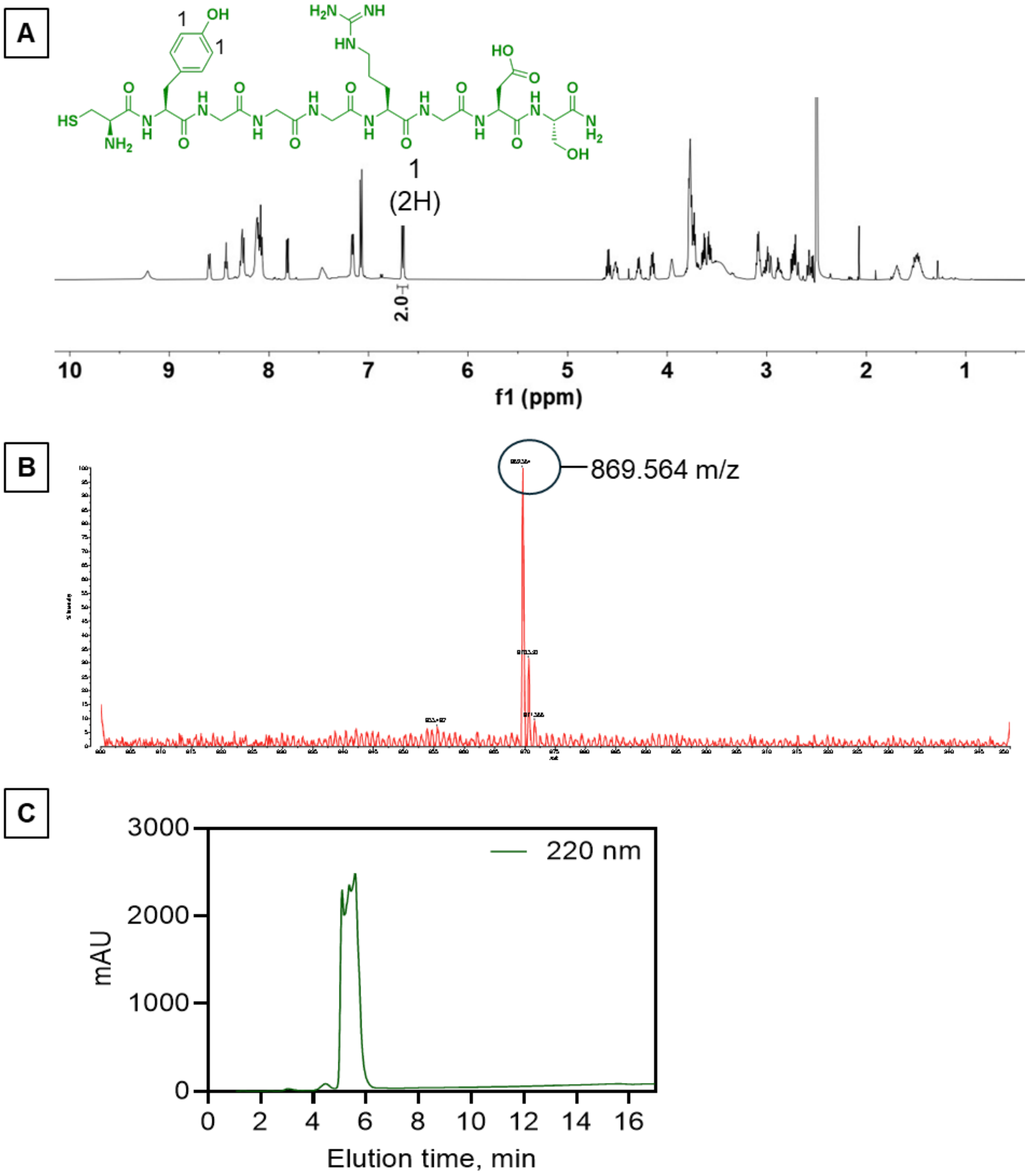

**Figure S6.** Representative $^{1}H$ NMR spectrum (500 MHz, DMF-$d_7$) and corresponding chemical structures of RGDS-PCL: δ = 6.77 (d, 4H, Ar-H, 1) and 4.07 (t, 456H, -O-$CH_2$-, a).

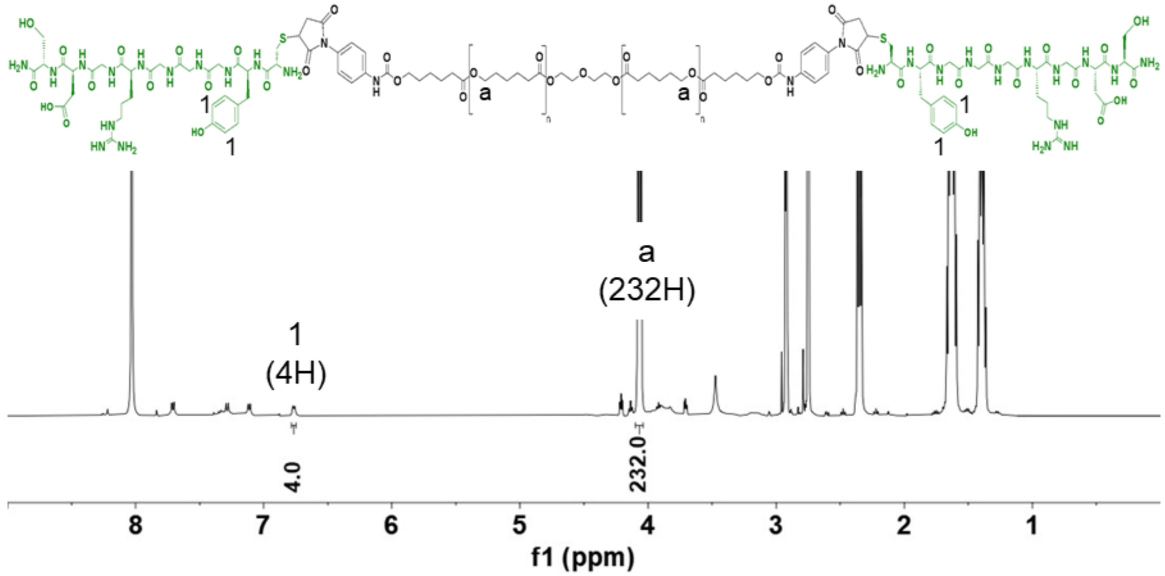
